# The microRNA processing subunit DGCR8 is required for a T cell-dependent germinal center response

**DOI:** 10.1101/2022.07.31.501995

**Authors:** Patrick Daum, Julia Meinzinger, Sebastian R. Schulz, Joana Côrte-Real, Manuela Hauke, Edith Roth, Wolfgang Schuh, Dirk Mielenz, Jürgen Wittmann, Hans-Martin Jäck, Katharina Pracht

## Abstract

We have previously shown that the microRNA (miRNA) processor complex consisting of the RNAse Drosha and the DiGeorge Critical Region (DGCR) 8 protein is essential for central B cell maturation. To determine whether miRNA processing is required to initiate T cell-mediated antibody responses, we deleted DGCR8 in maturing B-2 cells by crossing a mouse with loxP-flanked DGCR8 alleles with a CD23-Cre mouse. As expected, non-immunized mice showed reduced numbers of mature B-2 cells and IgG-secreting cells and diminished serum IgG titers. In accordance, germinal centers and antigen-specific IgG-secreting cells were absent in mice immunized with T cell-dependent antigens. Therefore, DGCR8 is required to mount an efficient T cell-dependent antibody response. However, DGCR8 deletion in B-1 cells was incomplete, which explains relatively unaffected B-1 cell numbers and adequate IgM and IgA titers in DGCR8-knock out mice and suggests that this mouse model could be used to analyze B-1 responses in the absence of functional B-2 cells.

## INTRODUCTION

MicroRNAs (miRNAs) are small non-coding single-stranded epigenetic regulators initially found to control the larva development of *Caenorhabditis elegans* (1). Higher eukaryotes use miRNAs to regulate gene expression at the post-transcriptional level (2), and altered miRNA expression is associated with many diseases (3,4).

Canonical miRNA maturation begins in the nucleus by synthesizing long primary transcripts (pri-miRNA). First, pri-mRNAs are processed to a ∼70 nucleotide long pre-miRNA duplex with a lariat structure by the heterotrimeric microprocessor complex consisting of the RNA-binding protein DGCR8 (*DiGeorge syndrome chromosomal/critical region 8*) and the RNAse III Drosha (5,6). Next, pre-miRNAs are exported into the cytoplasm by exportin-5, where they are further processed by the RNAse DICER1-TRBP complex to a short double-stranded RNA molecule (7). The leading strand of this duplex is integrated into the *RNA-induced silencing complex* (RISC). The mature miRNA then guides the RISC to its target mRNAs, which results in its degradation or inhibition of translation (8).

Studies analyzing the effect of targeted deletion of Dicer (9), DGCR8 (10), as well as members of the Argonaut-protein family (11) showed that these components of the miRNA processing machinery are essential in a variety of tissues. For example, B cell-specific ablation of Dicer or DGCR8 in early B cell precursors resulted in a developmental block at the pro-B cell stage (12,13). Furthermore, Dicer deficiency in germinal center (GC) B cells revealed an impairment of GC formation and T cell-dependent (TD) antibody responses (14). In contrast, the conditional CD19-Cre-mediated deletion of Dicer resulted in an altered B cell receptor (BCR) repertoire and high serum titers of auto-antibodies (15).

To investigate the role of DGCR8 and the canonical miRNA processing pathway in the establishment of the mature B-2 population consisting of follicular (FO) and marginal zone (MZ) B cells and their antigen-dependent activation, DGCR8 deletion was induced during the maturation process of splenic transitional B-2 cells by crossing a transgenic CD23-Cre mouse (16) to a mouse strain with loxP-flanked DGCR8 alleles (12). Here we demonstrate that CD23-Cre-mediated DGCR8 deletion in mice impaired the establishment of follicular (FO) B-2 cells and led to a reduction in IgG serum titers and the number of antigen-specific IgG-secreting cells. Furthermore, analysis of the cell viability of *in vitro* generated plasmablasts showed that ablation of mature miRNAs compromised the survival potential of LPS-activated mature B cells, which could, at least in part, mechanistically explain faulty TD-dependent antigen-specific humoral immune response with reduced GC formation.

## RESULTS

### B cell-specific DGCR8-deficiency reduces serum IgG titers and impairs the establishment of mature B cells as well as IgG-secreting cells *in vivo*

To investigate the role of DGCR8 in maturing B-2 cells, mice with loxP-flanked (floxed) DGCR8 alleles (12) were crossed to CD23-Cre transgenic mice (16). In this mouse line, the Cre activity is coupled to the expression of the FCepsilon receptor II (a) promoter, also known as CD23(a). Expression of CD23 is induced in secondary lymphatic organs at transitional stage 2 of immature B cells (T2- and T3-B cells) and in FO B cells (17). T2 B cells are the common precursor of marginal zone (MZ) and FO B cells.

To test DGCR8 deletion efficiency in mature B-2 cells, genomic DNA from splenic murine MZ (CD19^+^CD21^+^CD23^low^)- and FO (CD19^+^CD21^low^CD23^+^) B cells of CD23-Cre DGCR8^fl/fl^ mice (DGCR8-bKO) and CD23-Cre DGCR8^wt/w^ (Cre) littermates was PCR amplified with primers flanking the floxed exon 3 of the DGCR8 gene locus (Figure 1A). As expected, CD23-Cre-mediated DGCR8 deletion was efficient in FO and MZ B cells. Therefore, DGCR8-bKO mice were used to analyze the role of mature miRNAs in establishing the splenic B-2 cell populations and their TD-dependent antigen-driven activation.

**Figure 1:**
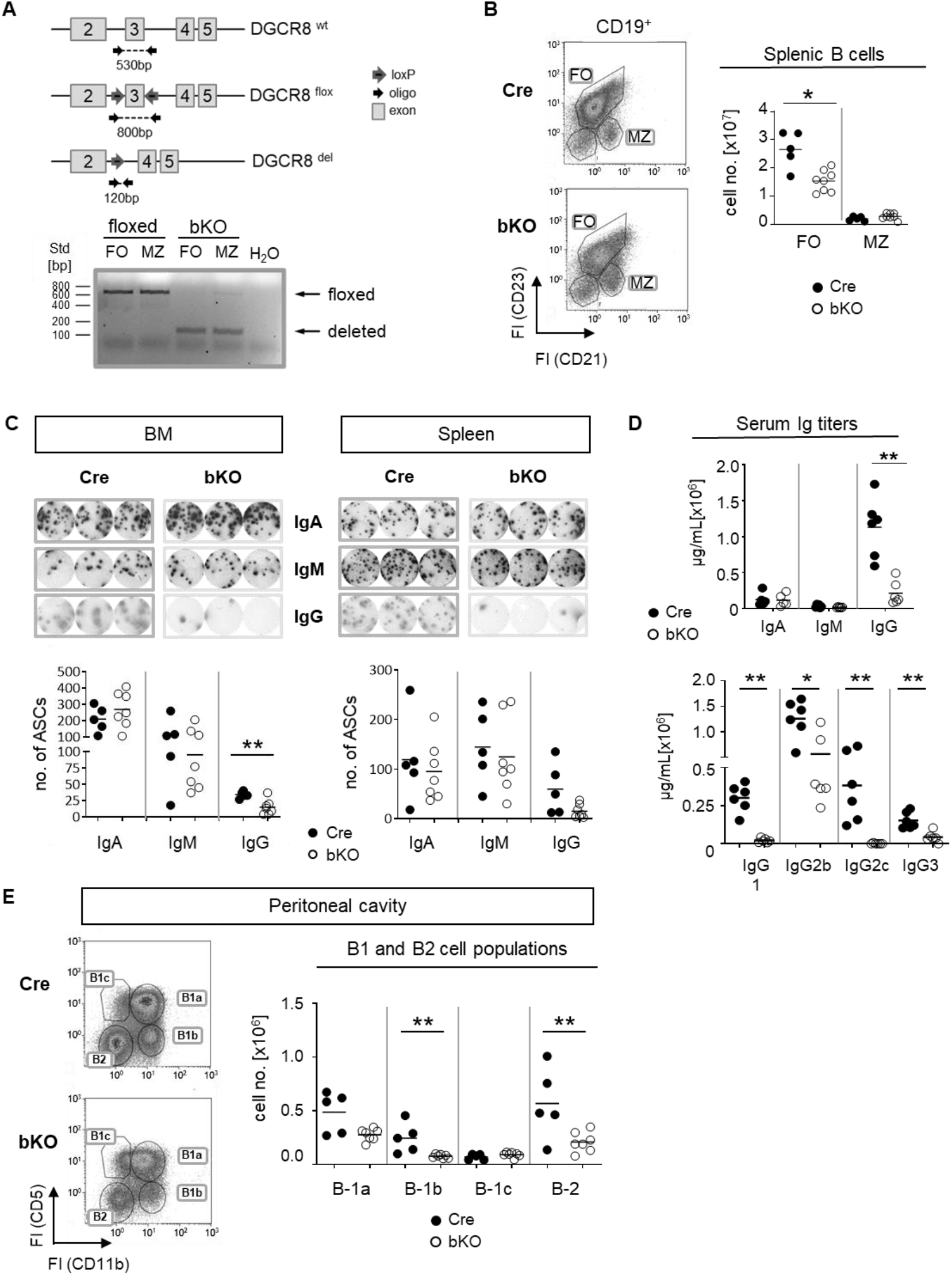
B cell-specific DGCR8 deficiency impairs the establishment of mature B cells and IgG-secreting cells *in vivo*. **A)** Construction of a mouse carrying a DGCR8 allele with a floxed exon 3 (upper panel). The locus was analyzed by PCR of flow cytometry-sorted MZ B cells (CD19^+^CD23^low^CD21^+^) and FO B cells (CD19^+^CD23^+^CD21^low^) from DGCR8-bKO and flox-only control mice (Cre), respectively. PCR products are ∼530 bp for the wildtype, ∼800 bp for the floxed and ∼120 bp long for the Cre-deleted DGCR8 allele. **B)** Flow cytometry analysis of splenic B cells from DGCR8-bKO (bKO) and Cre control mice. Stained samples were pre-gated on CD19-positive cells. Cell numbers of MZ- and FO B cells in the total spleen cell populations were quantified by flow cytometry using flow count beads. Bars indicate the median of n=5 (Cre) or n=8 (bKO) mice. **C)** ELISpot assay to quantify cells secreting IgA, IgM or IgG in the bone marrow (left panel) and the spleen (right panel) from non-immunized DGCR8-bKO and Cre control animals. Depicted wells show the results from a total of 33.333 seeded cells. **D)** Serum samples from non-immunized DGCR8-bKO and Cre control mice of 9-18 weeks in age were analyzed by ELISA for total IgA, IgM, IgG and IgG-subclasses. n=6 mice of each genotype. **E)** Flow cytometry analysis of samples from peritoneal lavages of DGCR8-bKO and Cre control mice, to quantify B-1a (CD19^+^CD5^+^CD11b^+^), B-1b (CD19^+^CD5^-^CD11b^+^), B-1c (CD19^+^CD5^+^CD11b^-^) and B-2 (CD19^+^CD5^-^CD11b^-^) cell populations. Cell numbers were calculated per total lavage. Bars indicate the median of n=5 (Cre) or n=7 (bKO) mice. Each dot represents one mouse. Mann-Whitney test was used for statistical analysis. **p<0.01; * p<0.05.

To analyze if DGCR8 loss affects the development of mature splenic B cell populations *in vivo*, we investigated non-immunized DGCR8-bKO and Cre control mice. Flow cytometry analysis of DGCR8-bKO mice revealed a significant decline (∼1.7 fold) of splenic FO B cell numbers in DGCR8-bKO mice. In contrast, the MZ B cell population was not significantly altered (Figure 1B).

Since FO B cells comprise the majority of the mature B-2 cell population and are mainly contributing to the generation of T cell-dependent antigen-specific humoral immunity (18), we speculated that the decreased number of FO B cells (Figure 1B) impacts the number of IgH class-switched antibody-secreting cells (ASC). As expected, ELISpot analyses revealed an apparent decrease in IgG-positive ASCs in bone marrow (BM, significant) and spleen (not significant) of non-immunized DGCR8-bKO mice compared to Cre-control mice (Figure 1C). However, numbers of IgM- and IgA-positive ASCs were not altered in both organs of DGCR8-bKO and Cre mice (Figure 1C).

This effect of DGCR8 ablation on the development of IgG-secreting cells (Figure 1C). was verified by Ig Elisa in sera from DGCR8-bKO mice, i.e., a severe reduction in serum IgG (∼4.9x fold decrease) was observed in bKO mice. In contrast, total serum IgM and IgA were unaltered (Figure 1D). Interestingly, the IgG1 and IgG2c subclasses were almost undetectable in DGCR-bKO mice. On the other hand, the subclasses IgG2b and IgG3 were also reduced but could still be detected in the serum of bKO mice.

IgG2b and IgG3 subclasses are secreted by *in vitro* stimulated B-1 cells (19,20). In addition, B-1a cells develop from CD23-negative precursors, are self-renewing and provide most of the natural IgM and half of the serum IgA (21). In support, the floxed DGCR8 allele could still be detected by DNA-PCR peritoneal B1a cells isolated from DGCRbKO mice (not shown). Therefore, it is tempting to speculate that most of the serum IgA and IgM and the detectable IgG subclasses in DGCR8-bKO mice are provided by plasma cells originating from CD23-negative B-1a cells (22). To address this hypothesis, we examined B-1 cell populations in the preferred location, i.e., the peritoneal cavity. As expected, flow cytometry analyses revealed that bone marrow-derived B-1b and B-2 cells were reduced by half in the peritoneal cavity of DGCR8-bKO animals. In contrast, the numbers of fetal liver-derived B-1a and B-1c cells were not significantly altered (Figure 1E). Hence, unaltered numbers B-1a population in the peritoneal cavity of DGCR8-bKO mice can provide serum IgM, IgA, and likely IgG2b/3 detected in non-immunized DGCR8 (Figure 1D). In addition, B-1a cells can also reside in the spleen and bone marrow (23), supporting the detection of IgM- and IgA-secreting cells in these tissues of DGCR8-bKO mice (Figure 1C).

### DGCR8 is essential for germinal center formation

To determine whether DGCR8 ablation in T2-originated B-2 cells affects the T cell-dependent (TD) germinal center (GC) reaction and, therefore, IgG serum titers and numbers of IgG-secreting ASC, we induced a robust TD-immune response by .injecting mice with sheep red blood cells (SRBC). Then, 7 days later, mice were sacrificed, and GC formation was analyzed. (Figure 2A). Flow cytometry analysis of splenic cells from DGCR8-bKO mice showed an apparent but non-significant reduction in the numbers of CD19^+^PNA^+^ GC B cells compared to Cre control animals (Figure 2B). GC B cells were further analyzed for expression of GL-7 and CD95, two additional proteins separating cycling GC B cells and more mature B cells already primed to differentiate into plasmablasts (24). Dividing GC B cells (CD95^+^GL-7^+^) were significantly reduced in DGCR8-bKO mice (11-fold), whereas the cell population preponed for plasmablast differentiation (CD95^+^GL-7^low^) was slightly but significantly diminished (1.4-fold). Histological analysis from the same spleens confirmed the inability of DGCR8-bKO mice to form morphologically adequate GCs (Figure 2C). Contrary to Cre control animals, DGCR8-bKO mice were not able to mount specific structures such as a GC dark- (PNA^+^) or light zone (PNA^+^Ki67^+^) within the mantle zone (IgD^+^) of the B cell follicle.

**Figure 2:**
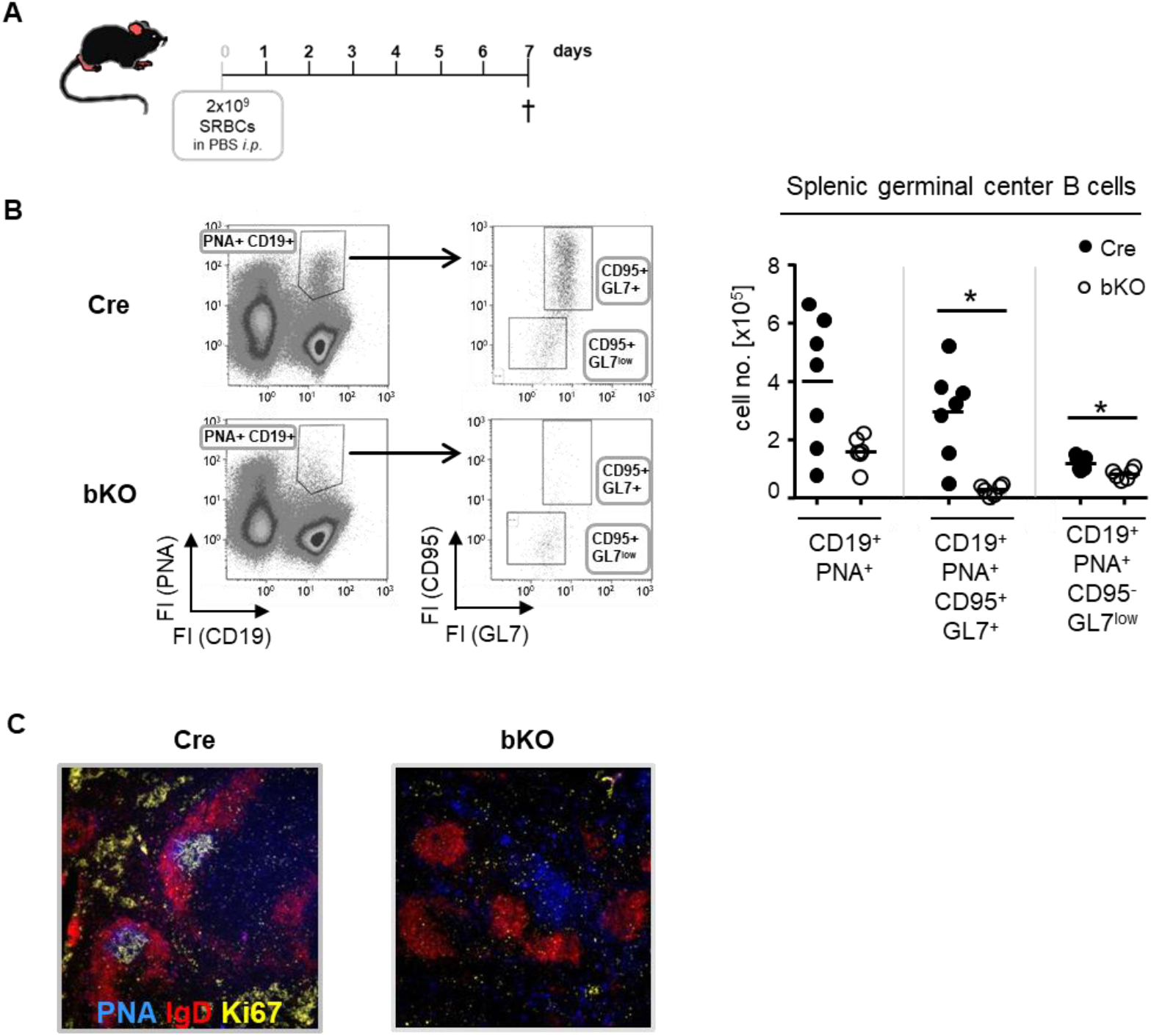
DGCR8 is essential for germinal center formation. **A)** DGCR8-bKO mice and Cre control animals were immunized with sheep red blood cell (SRBC) and analyzed one week later. **B)** Flow cytometry analysis of splenic cells from DGCR8-bKO mice and Cre control animals treated in A). GC B cells were defined as CD19^+^PNA^+^ and further subdivided into CD95^+^GL7^+^ and CD95^-^GL7^-^ cells. Cell numbers were calculated for the whole spleen. n=6-7 mice per genotype. **C)** Immunofluorescence microscopy of splenic cryosections from mice treated in (A). Sections were stained for IgD (red), PNA (blue), and Ki67 (yellow) to visualize the FO mantle zone (IgD^+^), as well as the light- and dark zone of the GC (PNA^+^ or Ki67^+^). Each dot represents a mouse and bars the mean of all mice with the same genotype. The Mann-Whitney test was used for statistical analysis. *p<0.05; ** p<0.01; *** p<0.001.

Based on these findings, we conclude that CD23-Cre-mediated ablation of DGCR8 in transitional B cells severely affects the establishment of GCs, which explains the reduced serum IgG titers and diminished numbers of IgG-secreting cells in DGCR8-bKO mice (Figure 1).

### DGCR8-bKO mice fail to mount a TD antigen-specific IgG response

We could demonstrate that DGCR8-bKO mice resemble a phenotype characteristic for hypogammaglobulinemia, with reduced basal levels of serum IgG, diminished numbers of IgG-secreting cells and disturbed formation of GC B cells and structures (Figure 1C, 1D and 2).

To determine whether DGCR8-bKO mice generate an antibody response to a TD antigen, mice were immunized with the antigen TNP-KLH in alum, boosted with TNP-KLH in PBS and finally analyzed on day 49 after primary immunization (Figure 3A). Strikingly, DGCR8-bKO mice completely failed to produce NP-specific IgG antibodies in response to a first immunization, and they responded just poorly, if at all, to a re-challenge with the same antigen (Figure 3B). In contrast, the production of TNP-specific IgM antibodies was not affected in DGCR8-bKO mice and was likely mounted by B-1 cells.

**Figure 3:**
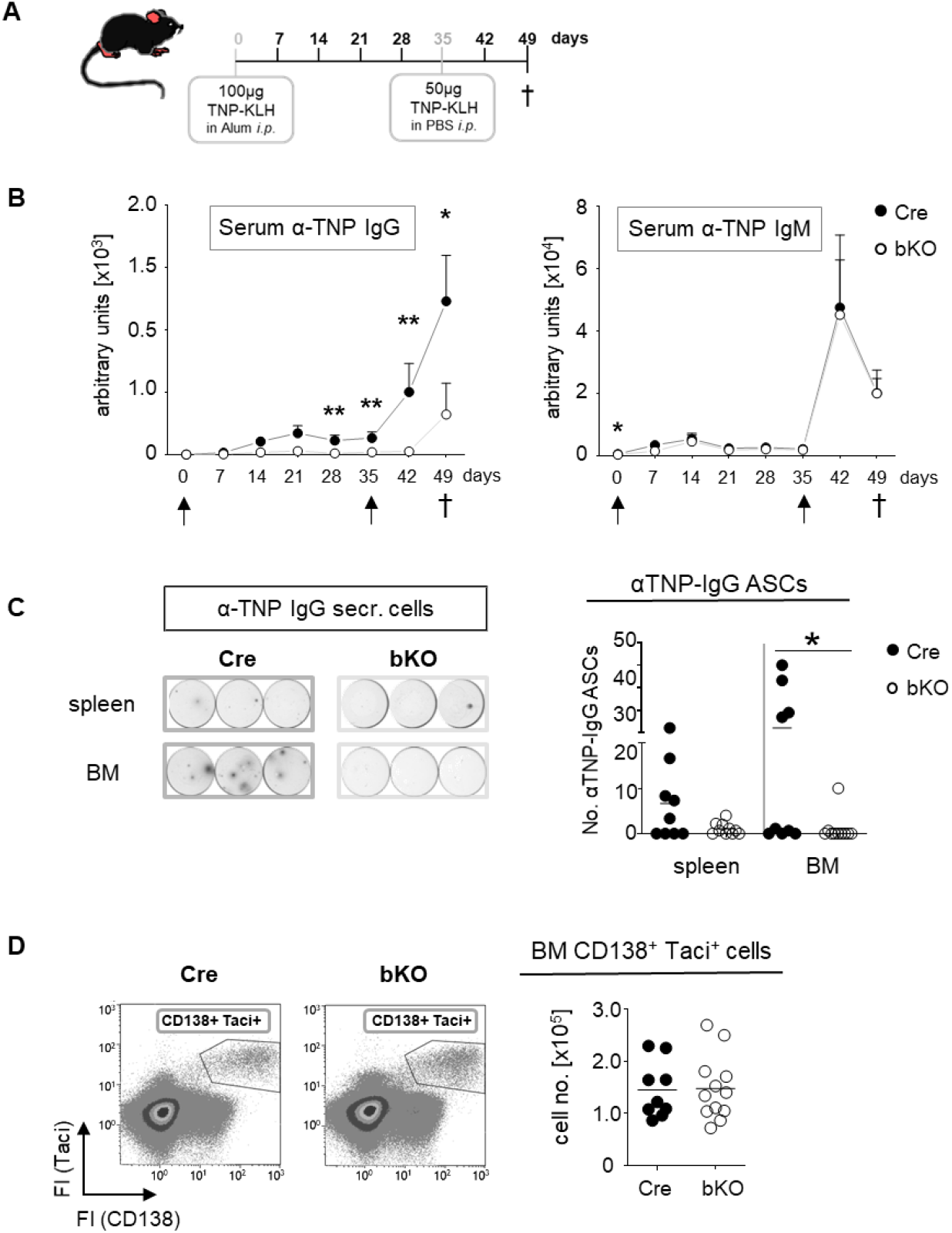
CD23-Cre-mediated DGCR8-deficient mice fail to mount a TD antigen-specific IgG response. **A)** DGCR8-bKO and Cre control mice were injected intraperitoneally with TNP-KLH (9-18 weeks in age) in alum, re-challenged on day 35 and sacrificed 49 days after primary immunization. **B)** Blood was collected weekly, and sera were analyzed for TNP-specific IgG and IgM by ELISA. Arrows indicate immunization time points. Each dot represents the mean (+SEM) of n=9 (Cre) or n=12 (bKO) mice. **C)** Mice were sacrificed on day 49 p.i. (post-immunization), spleen and bone marrow cell suspensions were analyzed for TNP-specific IgG secreting cells by ELISpot assay. The examples show the results obtained from 666.667 cells incubated per well. Cell numbers are calculated per 2×10^6^ seeded cells. **D)** Flow cytometry analysis of bone marrow cells. Plasma cells and plasmablasts were defined as CD138^+^ Taci^+^ cells (25). Cell numbers were calculated for one tibia and one femur. **C)** and **D)** Bars indicate the median of n=9 (Cre) or n=12 (bKO) mice from N=3 experiments. Each dot represents an individual mouse. The Mann-Whitney test was used for statistical analysis. *p<0.05; ** p<0.01; *** p<0.001.

ELISpot analysis showed a severe reduction of TNP-specific IgG-secreting cells in the bone marrow of DGCR8-bKO mice (Figure 3C), supporting the serum titer results in Figure 3B. Except for one individual mouse, no TNP-specific IgG-secreting cells were detectable in the bone marrow of DGCR8-bKO mice. A similar trend was observed in spleens of the same mice; even so, alterations were insignificant. However, the total numbers of CD138/Taci-positive plasmablasts and plasma cells (25) in the bone marrow of immunized DGCR8-bKO mice did not differ from those of Cre control animals (Figure 3D), which could be explained by the presence of a functional B-1a compartment in DGCR8 bKO mice.

In summary, B-2 cell-specific abrogation of DGCR8 by CD23-Cre is associated with a lack of antigen-specific serum IgG (Figure 1C and D) and the inability of the mice to generate or maintain IgG-secreting plasma cells upon TD immunization (Figure 3B and C). We, therefore, suggest that the two major hallmarks of GC reaction, namely IgH-isotype class switching and generation of high affine antibodies by somatic hypermutation, are disrupted in mice that lack expression of the miRNA processor component DGCR8 in mature B-2 cells.

### DGCR8-deficiency affects the viability of follicular and marginal zone B cells *in vitro*

A previous study from our lab has shown that the conditional deletion of DGCR8 in early B cell precursors blocked the central maturation of pro-B cells into mature B cells due to the impaired viability of pro-B cells (12). We, therefore, hypothesized that the inability of B-2 cells to form a GC reaction in DGCR8-bKO mice (Figure 2) could also be explained by a diminished survival potential of DGCR8-negative B cells before the onset of a GC reaction. To evaluate this idea, we loaded isolated splenic B cells from Cre-control and DGCR8-deficient mice with a proliferation fluorescence dye by following the rate of the decrease in the fluorescence intensities and determining the cell numbers of live cells (Figure 4A). As shown in Figure 4A, fluorescence-loaded DGCR8-deficient B cells showed a 2.9-fold higher fluorescence content of the used proliferation marker dye than the control cells, indicating retardation in cell proliferation.

**Figure 4:**
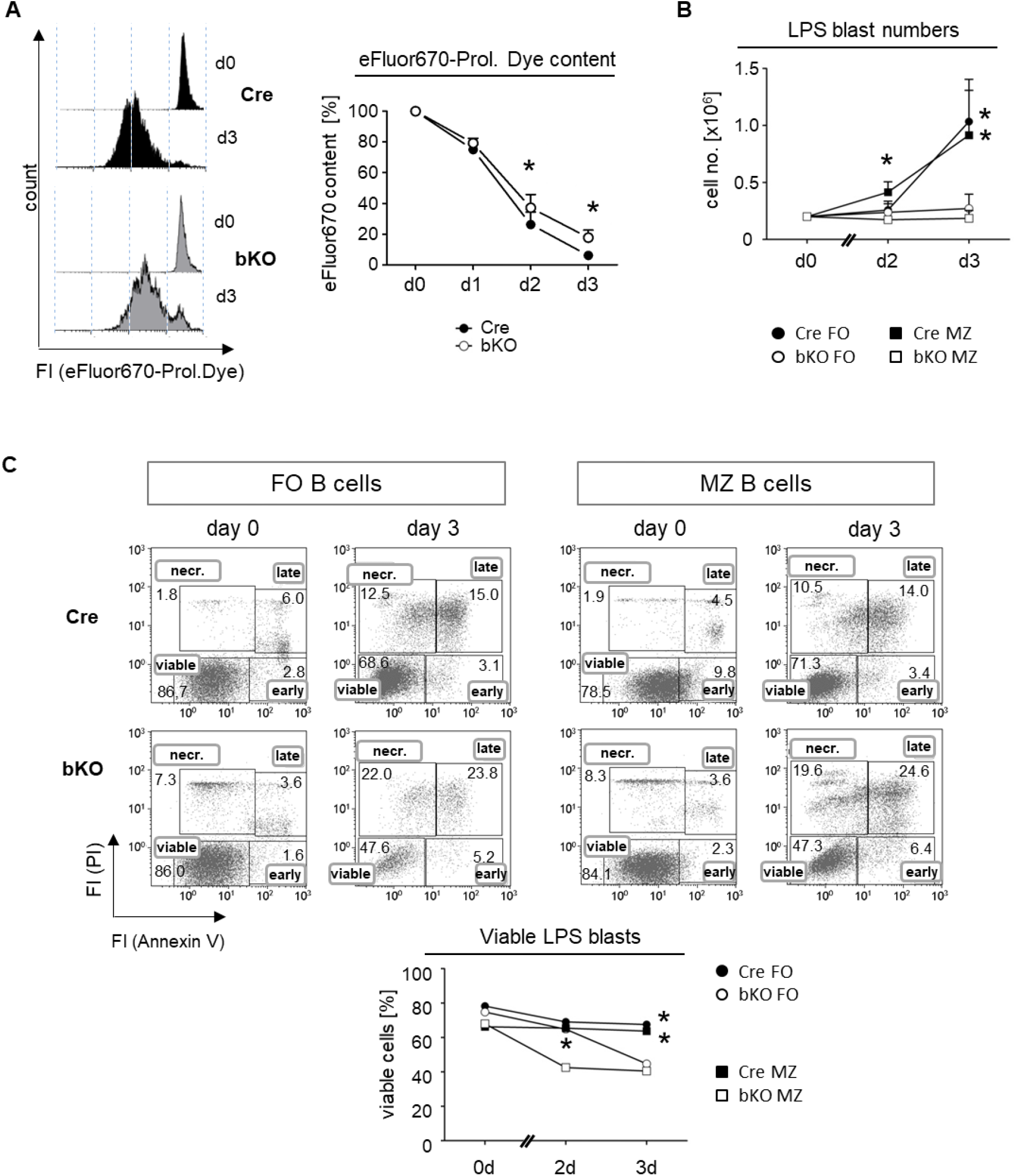
DGCR8-deficiency affects the viability of follicular and marginal zone B cells *in vitro*. **A)** Naive splenic B cells were isolated from Cre control and DGCR8-bKO mice by magnetic cell sorting (EasySep©) and analyzed for lipopolysaccharide (LPS)-induced proliferation *in vitro*. Cells were stained with the proliferation dye eFluor670 right after isolation, and the content of the dye was measured for three days using flow cytometry. The representative figure shows eFluor670 content (day 0 and day 3). The Graph depicts changes in eFluor670 fluorescence intensity of Cre and DGCR8-bKO cells normalized to the basic value determined on day 0. Points represent the mean from n=4 mice per genotype. **B)** Flow fluorescence-based flow cytometry-sorted splenic MZ B cells (CD19^+^CD23^low^CD21^+^) and FO B cells (CD19^+^CD23^+^CD21^low^) from Cre controls and DGCR8-bKO mice were stimulated *in vitro* with LPS. Cell numbers were determined using flow count fluorophores. Points represent the mean from n=3 mice per genotype. **C)** Cell viability in the samples described in B) was analyzed at different time points by flow cytometry with propidium iodide (PI) and AnnexinV. AnnexinV- and PI-negative cells were defined as viable. Mann-Whitney test was used for statistical analysis. * p<0.05; ** p<0.01; *** p<0.001.

Next, we isolated splenic FO and MZ B cells by fluorescence-activated cell sorting (FACS) and analyzed them *in vitro* for their viability under the same stimulatory conditions as described in Figure 4A by an AnnexinV/propidium iodide (PI)-assay. First, we observed that LPS activation resulted in the expansion of cell numbers in cultures of DGCR8-deficient FO B cells and MZ B cells. In contrast, B cells from Cre control animals expanded as expected in response to LPS (Figure 4B). Furthermore, the viability of the isolated DGCR8-deficient FO and MZ B cells decreased over time, showing 20% fewer AnnexinV/PI-negative viable cells compared to Cre controls after three days in culture (Figure 4C).

These results imply that DGCR8 ablation slightly but significantly impaired B cell proliferation upon stimulation but had an even more significant impact on cell viability. This explains the dramatic restriction in activated B cell expansion *in vitro*. In addition, the diminished survival potential of B cells upon antigen activation could also explain the defect in GC formation (Figure 3) and the inability to mount a high specific IgG-response (Figure 4) in immunized DGCR8-bKO mice.

## DISCUSSION

In this study, we demonstrate in mice with a B cell-specific conditional DGCR8 deficiency (DGCR8-bKO) that the ablation of microRNAs in maturing B-2 cells affects the humoral immune response before the onset of GCs drastically

The decreased cell viability of DGCR8-deficient FO B cells *in vitro* was in line with the observation of a diminished FO B cell population in the spleens of DGCR8-bKO mice. This result perfectly fits the reduced survival rate of Mb1-Cre-mediated DGCR8-deficient pro-B cells (12). Although isolated splenic DGCR8-deficient MZ B cells exhibit a defective survival upon stimulation *in vitro*, this population was not altered in cell numbers *in vivo*. Therefore, we assume that residual DGCR8 mRNA and mature miRNAs in MZ B cells might temporally bypass the deletion of the DGCR8 allele and mask a potential phenotype *in vivo* (Supplementary Figure 1). In addition, intercellular interactions like the secretion of B cell-activating factor (BAFF) by various lymphatic cells could temporarily suppress apoptotic processes in DGCR8-deficient B cells *in vivo* (26).

Nevertheless, analysis of B cell populations in non-immunized DGCR8-bKO mice implies that DGCR8 ablation in maturing B cells has a functional effect on forming antibody-secreting cells. In particular, the class switching to IgG-secreting cells and thereby the production of IgG is affected in DGCR8-bKO mice. Flow cytometry and fluorescence microscopy revealed that B cells could not form morphologically adequate GCs without mature miRNAs. Surprisingly, neither the class switching to IgA nor the formation of IgA- or IgM-secreting cells was significantly compromised in DGCR8-deficient mice. In addition, IgG2b secretion was slightly reduced, while other IgG-subclasses were absent in the serum of DGCR8-bKO mice. This particular phenotype is of most significant interest, as B-1 cells can switch to Ig-secreting cells of those IgH subclasses that were not changed in DGCR8-bKO mice (19–21) and numbers of CD23-negative self-renewable B-1a cells in the peritoneal cavity, originating from fetal liver cells, were expectantly unaffected (22). The presence of a functional B1 compartment was supported by our findings that DGCR8 deletion was incomplete in isolated B1a cells (Supplemental Figure #). Therefore, the humoral immune response observed in DGCR8-bKO mice is likely mounted by B-1 cells.

We found that IgA and IgM are the predominant IgH isotypes in non-immunized mice kept in our local animal facility. At the same time, IgG-secreting cells only represent a minor fraction of the total plasma cell population in the BM (Figure 1C). Accordingly, due to their low frequencies, the loss of IgG-secreting cells in the BM of DGCR8-bKO mice very likely does not affect the total number of plasma cells, which explains the unaltered numbers of BM CD138^+^Taci^+^ plasmablast/plasma cells in DGCR8-bKO mice compared to Cre-controls. Furthermore, only small numbers of plasma cells are usually found in the bone marrow (27); consequently, niches could be populated by non-IgG-secreting plasma cells that originated from B-1 cells in DGCR8-bKO mice.

Taking into consideration that DGCR8-bKO mice did not exhibit an adequate GC formation upon activation with TD-antigen, it is surprising that these mice can mount an antigen-specific and IgM memory-like response (against TNP), as documented in Figure 3B. However, several studies reported that memory B cells could originate from a GC-independent pathway early during an immune response (28–30). Furthermore, most IgM+ memory-like B cells are likely generated in a GC-independent manner (31, 32). Furthermore, as hypothesized before, most of the serum IgM and IgA detected in DGCR8-bKO mice could originate from B-1 as these cells can undergo TD-independent Ig class switch comparable to B-2 cells (33,34). Therefore, these findings partly explain the existence of the antigen-specific IgM memory B cells and the induction antigen-specific IgM, which we observed in DGCR8-bKO mice upon re-challenge with the same TD-antigen.

Our study clearly showed that the deletion of DGCR8 in maturing B2 precursors almost completely abolished a TD GC reaction. As the CD23-Cre-mediated deletion of DGCR8 is incomplete in B1a cells, this B cell compartment might be responsible for most of the antibody responses observed in DGCR8-bKO mice. Therefore, the CD23-Cre/DGCR8 KO mouse could be a unique and excellent model to study B1a responses and tumorigenic events without a functional FO and MZ B cell compartment.

## MATERIAL AND METHODS

### Mice

All mice were maintained under pathogen-free conditions in the Nikolaus-Fiebiger Center animal facility of the University of Erlangen-Nürnberg, Erlangen, Germany. All animal experiments were performed according to institutional and national guidelines. Transgenic CD23-Cre mice (Kwon et al., 2008) were crossed with loxP-flanked DGCR8-mice (Brandl et al., 2016). The mice have a C57Bl/6 background.

### Immunization of mice

Mice were immunized with 100 μg (100 μl in PBS) TNP-KLH (load 18, LGC Biosearch Technologies) in 100 μl alum (Imject Alum Adjuvants, ThermoFisher) intraperitoneally. On day 35, mice were boosted with 50 μg (50 μl) TNP-KLH in 50 μl alum intraperitoneally. Alternatively, mice were immunized with 2×10^9^ sheep red blood cells (SRBC, Fiebig Nähstofftechnik) in 300 μl PBS intraperitoneally.

### Flow cytometry analysis

1-2×106 isolated cells were stained in 96-well plates for flow cytometric analysis. Unspecific bindings were blocked by incubation with an unlabeled αCD16/32-antibody (eBioscience) for 15 minutes on ice. Afterward, surface markers were stained with the respective primary and secondary antibodies for 15-20 minutes on ice and in the dark. The AnnexinV/PI-staining was performed using the “Annexin Apoptosis Detection Kit APC “(eBioscience) following the manufacturer’s protocol with minor alterations (25μl Staining solution with AnnexinV-APC 1:100 per sample; PI 1:200). Proliferation analysis was performed using the Proliferation Dye eFluor™ 670 (eBioscience) according to the manufacturer’s protocol. Stained cells were acquired using a Gallios flow cytometer (Beckman Coulter). Raw data were analyzed using Kaluza (Beckman Coulter, Krefeld, Germany) software. The following antibodies were used for flow cytometric stainings: From BD Pharmingen αCD19 APC-Cy7 and αCD5 PE; from Vector PNA FITC; from BioLegend αCD138 Brilliant Violet 421; from eBioscience αCD11b FITC, αCD95 PE, αGL7 eFluor660, αTACI/CD267 PE, αCD23 FITC and αCD21 biotinylated; from Jackson ImmunoResearch Streptavidin Cy5.

### Magnetic- or flow cytometric B cell isolation and *in vitro* culture

To isolate splenic follicular or marginal zone B cells by FACS, cell suspensions were stained as described for flow cytometric analysis in PBS-2% FCS. Cells were stained in 15 ml reaction tubes. The solutions’ volumes were adjusted according to the used cell numbers. Cells were stained with αCD19 Brilliant Violet 421 (BioLegend), αCD23 PE (BioLegend), αCD21 biotinylated (eBiosciene) and Streptavidin Cy5 (ImmunoResearch) and isolated with a purity >99% with the MoFlo cell sorter (Beckman Coulter). Splenic naïve B cells were isolated by magnetic cell sorting using the “EasySep™ Mouse B Cell Isolation Kit” from STEMCELL according to the manufacturer’s protocol. Isolated splenic B cells were cultured in complete RPMI1640 with 10% FCS and 10 μg/mL LPS with a density of 2×10^5^ cells/mL (37°C, 5 % CO2).

### PCR

To analyze the DGCR8-alleles by PCR, genomic DNA was isolated from biopsies by proteinase K (Peqlab peqGOLD)-digestion in PBND-buffer and used the following gene-specific primers in a PCR: 5’-GATATGTCTAGCACCAAAGAACTCC-3’ and 5’-GATCTCAGTAGAAAGTTTGGCTAAC-3’. For the loxP-flanked exon 3, a fragment with 730bp is expected, for the wildtype allele a 500bp fragment, and for the deleted allele a 120bp fragment.

### RNA isolation

RNA, including microRNAs, were isolated using the “miRNeasy Mini Kit” (cell numbers ≥ 5 × 105, Qiagen, Cat# 217004). FACS-isolated FO or MZ B cells were directly sorted into the Qiazol Lysis Reagent (700 μl final volume). The samples were stored at −70°C until further processing. Thawed samples were vortexed for 1 minute and incubated at RT for 5 minutes before further processing following the manufacturer’s manual. RNA concentrations and purity (absorption at 260 nm and a ratio of 260/280 of ∼2.0, respectively) were determined using the NanoDrop ND-1000 (Peqlab).

### cDNA synthesis and TaqMan© qRT-PCR

Isolated miRNAs were transcribed in PCR templates (cDNA) using the “TaqMan MicroRNA Reverse Transcription Kit” (Applied Biosystems, Cat# 4366597) after digestion of the remaining genomic DNA by incubation with the “DNase I Kit” (Sigma) following the manufacturer’s manual. 5 μl RNA (2ng/μg) and 3 μl of 5x Primer stock (ThermoFisher Scientific, miR-29a-3p: Cat# 002112; miR-16-1: Cat# 000391 or RNU6B Cat# 001093) were added to the 7 μl master mix (0.15 μl dNTP, 1 μl transcriptase, 1.5 μl buffer, 0.2 μl RNase inhibitor and 4.15 μl RNAase-free water) for conversion of mature microRNAs to cDNAs. Mixtures were incubated in a PCR machine for 30 minutes at 16°C, 30 minutes at 42°C and 5 minutes at 85°C. The cDNA preparation was then pre-diluted 1:5 with RNAase-free water and quantified using the “TaqMan© qPCR analysis TaqMan Universal Master Mix II” (Invitrogen, Cat# 4427788). For each reaction, 5 μl cDNA (1:5), 0.75 μl miRNA-specific probe mix (20x), 7.5 μl master mix, and 1.75 μl RNAse-free water were mixed in 96-well plates (Thermo Scientific, Cat# AB-1100) and covered with “adhesive qPCR Plate Seals” (Thermo Scientific, Cat# AB-1170). TaqMan© qRT-PCR analysis was performed in the “7300 Real-Time PCR System” (Applied Biosystems). To detect DGCR8 mRNAs by SYBR Green PCR, mRNAs were converted to cDNAs using the “RevertAid First Strand cDNA Synthesis Kit” (Fermentas) following the manufacturer’s manual. Specific primers were designed using GETPrime (DGCR8 forward primer: AAGAATAAAGCTGCCCGAG; DGCR8 reverse primer: GTCTTTAGGCTTCTCCTCAG) (35). SYBR Green RT-PCR was performed using 7.5 μl of the SYBR Green PCR Master Mix (Thermo Fisher Cat# 4309155) and 1 μl cDNA (1:5 pre-diluted in ultrapure water). Each sample was measured in triplicates. Reactions without the cDNA template (NTC) served as a negative control to validate the specificity of the reaction. The mean of the Ct-values (cycle threshold) was calculated for the triplicates of each sample. The mean-Ct of the housekeeping gene RNU6B was subtracted from the mean-Ct of the respective microRNA, while β-Actin mRNA (Forward primer: TGGAATCCTGTGGCATCCATGAAAC; reverse primer: TAAAACGCAGCTCAGTAACAGTCC) served as a housekeeper gene for mRNA quantification (36). This value was used to calculate the ΔCt-values.

### ELISpot and ELISA

To identify the frequencies of antibody-secreting cells in the single-cell suspensions from the spleen or the bone marrow of mice, ELISpot analysis was performed in 96-well flat-bottom plates as described in (37). To analyze total Ig-secreting cells, the plates were coated with goat-α-mouse IgM, IgG or IgA (Southern Biotech), while TNP-BSA (load 5; LGC BioSearch Technologies) was used for the identification of TNP-specific antibody-secreting cells. Cell suspensions were incubated overnight. Alkaline phosphatase (AP)-coupled goat α-mouse IgG, IgM or IgA antibodies (Southern Biotech) were used as detection antibodies. 5-Bromo-4-chloro-3-indolyl phosphate p-toluidine salt (BCIP; SigmaAldrich) was used in ESA substrate buffer for detection. Spots representing single antibody-secreting cells were counted using the Immuno-SpotR© Series 6 Ultra-V Analyzer from C.T.L. and analyzed with the C.T.L. Software BioSpotR© ImmunoSpot 5.1.36. For detecting serum Ig by ELISA, 96-well plates were coated as described for ELISpot-analysis. As detection antibodies, either AP-coupled α-mouse-Ig (IgG, IgA or IgM) antibodies or HRP-coupled α-mouse-Ig (IgG, IgA or IgM) antibodies (Sothern Biotech) were used. For AP- and HRP-coupled detection antibodies, alkaline phosphatase yellow (pNPP) liquid substrate (Sigma) and TMB Substrate Reagent Set (BD Pharmingen) were used, respectively. ELISA plates were measured using SpectraMax 190 at 450 nm (HRP) or 405 nm (AP).

### Immune histology

For immune histological analysis, splenic tissue samples were frozen at -80°C in Tissue-Tek© O.C.T. © (Sakura), and sections were generated at the Leica CM3050S cryostat. Then, spleen sections were fixed in acetone (-20°C) and stained with the respective primary and secondary antibodies or chemicals. For the analysis of GCs, PNA Rhodamine (Vector), αIgD FITC (SouthernBiotech) and αKi67 APC (BioLegend) were used before the sealing with VectaShield (Vector).

### Statistical analysis

Significances and p-values were determined using the GraphPad Prism software (GraphPad Software, La Jolla, CA, USA). Statistical tests were performed as indicated below each figure.

## ABBREVIATIONS

ASC=: antibody-secreting cell;
BCR=: B cell receptor;
BM=: bone marrow;
DGCR8: DiGeorge Critical Region 8;
FO=: follicular;
GC=: germinal center;
Ig=: immunoglobulin;
miRNA=: microRNA;
MZ=: marginal zone;
PI=: propidium iodide;
RISC=: RNA induced silencing complex;
SRBC=: sheep red blood cells;
TD=: thymus-dependent;
TNP-KLH=: 2,4,6-Trinitrophenyl Keyhole Limpet Hemocyanin.

## ACKNOWLEDGMENTS

This work was supported in part by research grants GRK1660 (to HMJ and DM) and TRR130 (to HMJ, DM and WS) from the Deutsche Forschungsgemeinschaft (DFG). JCR is a fellow of the Alexander-von-Humboldt Foundation and was supported by the program to promote equal opportunities for women in research and teaching (FFL) from the FAU Erlangen-Nürnberg. We thank Uwe Appelt and Markus Mrotz for MoFlo cell sorting.

## AUTHOR CONTRIBUTIONS

PD, KP, JM, JCR, MH, SRS and ER performed experiments. PD and HMJ designed experiments. PD and KP analyzed and visualized the data. PD, KP and HMJ interpreted the data. JM, WS, DM, SRS and JW provided scientific input for data interpretation. HMJ and JW conceptualized the project. HMJ supervised the project. PD, KP and HMJ wrote the manuscript.

## MATERIALS & CORRESPONDENCE

Correspondence and material requests should be directed to the lead contact, Hans-Martin Jäck.

## SUPPLEMENTAL INFORMATION

### SUPPLEMENTAL FIGURES

**Supplementary Figure 1:**
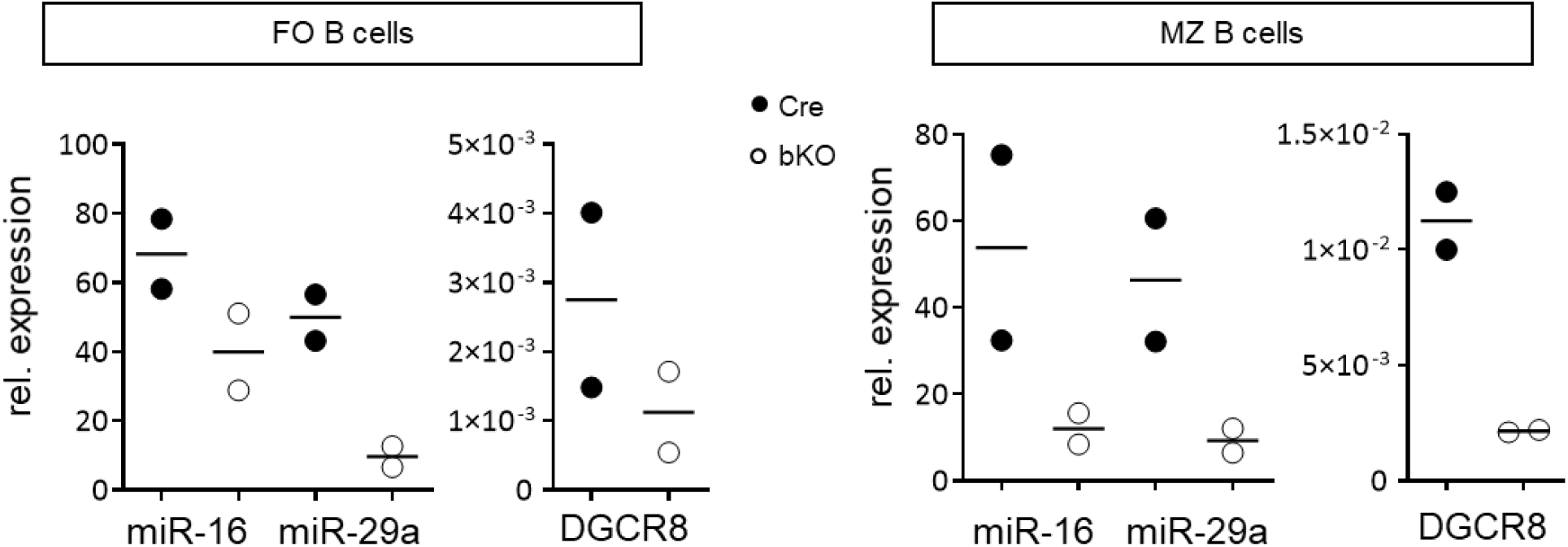
Deletion efficiency in splenic cells of DGCR8-bKO mice. A) Splenic follicular (FO: CD19^+^CD23^+^CD21^+^) and marginal zone (MZ: CD19^+^CD23^-^CD21^+^) B cells of DGCR8-bKO mice and Cre control animals were isolated by FACS. microRNAs and mRNAs were isolated, reverse transcribed to cDNA and analyzed by TaqMan© qPCR (miRNA) or SYBR Green RT-PCR (mRNA). CT values of the housekeeping genes RNU6B (miRNA) or β-Actin (mRNA) were used to calculate ΔCT values. n=2

